# Release of fragmented host, cell-free, genomic DNA into the circulation of pigs during infection by virulent African swine fever virus

**DOI:** 10.1101/2023.09.06.556476

**Authors:** Ann Sofie Olesen, Louise Lohse, Camille Melissa Johnston, Thomas Bruun Rasmussen, Anette Bøtner, Graham J. Belsham

## Abstract

African swine fever virus (ASFV) causes a severe hemorrhagic disease in domestic pigs resulting in high case fatality rates. The virus replicates in circulating cells of the monocyte-macrophage lineage and within lymphoid tissues, e.g. tonsils, spleen and lymph nodes. The infection results in high fever and a variety of clinical signs from about 3 days post infection. In this study, it was observed that one of many changes resulting from ASFV-infection within pigs was a large (>1000-fold) increase in the level of circulating cell-free DNA (cfDNA), including the beta-actin gene, derived from the nuclei of host cells, in the serum. This change occurred in parallel with the increase in circulating ASFV DNA. In addition, elevated levels (about 30-fold higher) of host mitochondrial DNA (mtDNA) were detected in serum from ASFV-infected pigs, but with a much higher baseline level of mtDNA in sera from uninfected pigs. The host derived cfDNA is derived from dead cells which may, or may not, have been infected. For comparison, the release of the cellular enzyme, lactate dehydrogenase (LDH), a commonly used marker of cellular damage, was also found to be elevated during the infection. The cfDNA is readily detected in serum and is a more sensitive host marker of ASFV infection than the release of mtDNA or LDH. In addition, sera from pigs infected by classical swine fever virus (CSFV), which causes a clinically similar disease as ASFV, were also tested but this infection did not result in the release of cfDNA, mtDNA or LDH.

**Author summary:** African swine fever virus causes a severe hemorrhagic disease in domestic pigs and wild boar, which often leads to death within a week. The infection results in a spectrum of different clinical signs and other changes within infected animals. In this study, we have shown, for the first time, that one consequence of infection by a highly virulent strain of this virus is the release into the blood of host genomic DNA, in a highly fragmented form. We found an increase of >1000-fold in the level of this cell-free DNA within the serum of infected animals. Furthermore, we also showed that the level of the small circular DNA from the cell mitochondria is also elevated in serum from infected animals as is the cellular enzyme lactate dehydrogenase but these changes were less marked and occurred later. The increase in the level of the cell-free host DNA is coincident with the increase in level of the viral DNA within blood and may act as a marker for infection by a highly virulent form of the virus. Remarkably, pigs infected by classical swine fever virus, which produces similar clinical signs, did not have elevated levels of these markers in their serum.

## Introduction

African swine fever (ASF) is a severe hemorrhagic disease of domestic pigs and other members of the family *Suidae*, including wild boar [1, 2]. The disease is caused by infection with African swine fever virus (ASFV) and can have a case fatality rate of up to 100% in both domestic pigs and wild boar. In consequence, the disease can cause major economic losses as well as serious animal welfare issues. The virus is classified within the genus *Asfivirus*; it is a large DNA virus belonging to the *Asfarviridae* family, indeed it is the only member of this family [3]. The viral genome is about 190 kbp in length and includes over 150 genes, many of which have unknown functions [4].

There are over 20 different genotypes of the virus (identified from the sequence of the gene encoding VP72), which have been identified in various locations across Africa [5]. Different ASFV strains can vary markedly in their virulence. A highly virulent, genotype II, virus has become important globally following its introduction into Georgia in 2007 and its subsequent spread into many countries in Europe and Asia [6]. Recently, the disease has also occurred in Haiti and The Dominican Republic [7]. It has been introduced into multiple EU countries and in 2022 new outbreaks occurred in eight of them including Germany, Italy, Slovakia and Poland [8]. In 2018, the virus spread further into Asia and has caused massive losses within the pig production industry in China and nearby countries, e.g. Vietnam, Laos, Cambodia and South Korea [9]. The virus continues to cause outbreaks in these different regions.

Infection of domestic pigs results in high fever (often >41 °C) together with a range of rather non-specific clinical signs, including lethargy and anorexia, that occur within a few days of infection [10, 11]. The presence of skin hemorrhages is often observed [12, 13], while vomiting and bloody diarrhea are sometimes recorded [12, 14]. Death can occur with few external signs but, during post mortem examinations, enlargement of the spleen and lymph nodes is typically observed along with internal bleeding [12, 14, 15]. The virus replicates in macrophages and monocytes within the blood but is also present at high levels within the tonsils, other lymph nodes, and the spleens of infected animals [2, 16, 17]. A feature of ASFV-infection in pigs is the loss (through apoptosis) of B- and T-lymphocytes in lymphoid tissues linked to the presence of infected monocytes [18]. It has been suggested that the infected monocytes signal to uninfected lymphocytes to enter apoptosis [19]. However, cell death by necrosis may also occur [11]. Hence, pigs that are acutely infected with ASFV can display severe lymphopenia [10, 20].

The death of nucleated cells can result in the release of cellular, genomic, DNA into the circulation system. This cell-free DNA (cfDNA) can be used as a biomarker for cell damage during organ transplant rejection or as a marker for various cancers (see review [21]). The concentration of cfDNA in plasma is normally very low but can be higher in serum due to some lysis of cells during blood clotting. The process of cell death by apoptosis results in the release of cellular contents within a range of extracellular vesicles, these can contain a variety of different molecules, including DNA [22]. These vesicles are present within serum/plasma.

The presence of the fragmented cfDNA in serum/plasma can be readily detected using sensitive real-time quantitative PCR (qPCR) assays that have a small target size.

In this study, the production of cfDNA, derived from the host genome, in the blood of ASFV-infected pigs has been examined in parallel with other markers of cellular damage. It appears that the production and release of cfDNA, derived from the host genome, into the serum is closely linked to the replication of ASFV in the infected pigs. For comparison, sera from pigs infected with low and high virulence strains of classical swine fever virus (CSFV, a pestivirus), which can cause similar clinical signs of disease and severe lymphopenia (with highly virulent strains) in pigs as with ASFV, were also assayed for the same markers.

## Materials and Methods

### Samples from ASFV-infected pigs

#### Experiment A

From an experiment performed in 2022 (here termed experiment A), the samples used here were from 4 male pigs (Landrace x Large White), about 12 weeks of age (numbered 13, 14, 15 and 20), which had been inoculated, using the intranasal route, with 4 log_10_ TCID_50_ of the genotype II ASFV POL/2015/Podlaskie strain (as used previously [23]). EDTA-blood samples were obtained from these pigs at 0 dpi, 3, 5, 6 and 7 (euthanasia) dpi, while serum samples were obtained at 0 dpi and at euthanasia only. Some separate results from this experiment, have been described previously but there is no overlap with the samples analysed here [24].

#### Experiment B

In another experiment, from 2020 (here termed experiment B), 12 male pigs (Landrace x Large White) were inoculated by the intranasal route with 4 log_10_ TCID_50_ of the ASFV/POL/2015/Podlaskie, as above. Results from analysis of certain samples (EDTA-stabilized blood (EDTA-blood) and peripheral blood mononuclear cells (PBMCs)) from these pigs have been described previously [23, 25] but no analysis of serum samples, which are described here, has been reported previously. The serum samples were obtained from blood samples collected prior to inoculation at 0 dpi and at 3, 5 and 6 dpi. The pigs were euthanized at 6 dpi.

In both experiments, water and a commercial diet for weaned pigs were provided *ad libitum*. EDTA-blood and unstabilized blood samples (for serum preparation) were collected prior to inoculation on day 0 and at indicated days post inoculation (dpi). All samples were stored at -80◦C until further analysis. Rectal temperatures were recorded and a total clinical score was calculated on all sampling days, as described previously [23]. The pigs were euthanized at the end of the study period by intravascular injection of Pentobarbital following deep anesthesia.

Animal care and maintenance, experimental procedures and euthanasia were conducted in accordance with EU legislation on animal experimentation (EU Directive 2010/63/EU). The original animal experiments were approved by the Ethical and Animal Welfare Committee of the Generalitat de Catalunya (Autonomous Government of Catalonia; permit number: CEA- OH/11744/2) and no new animal experiments were performed for the analyses presented here.

### Samples from CSFV-infected pigs

Serum samples had been collected from six pigs that had been inoculated with a high virulence genotype 2.1 strain (CSF1047, Israel, 2009) or a low virulence genotype 2.2 strain (CSF0906, Bergen); this study has been described previously [26]. Briefly, the pigs, which originated from a standard Danish swine herd, were inoculated by the intranasal route with 5 log_10_ TCID_50_ of the CSFV (three pigs for each virus strain). Serum used for the current study was prepared from blood samples collected prior to inoculation at 0 days post infection (dpi), and at 4, 7, 10, 11 or 22 dpi as indicated [26]. These serum samples were stored frozen at -20 ◦C until further analysis.

### Laboratory analyses

#### Viral genome detection

Nucleic acids were purified from whole blood or serum using the MagNA Pure 96 system (Roche) with the DNA/Viral NA 2.0 kit and the Viral NA Plasma external lysis S.V. 3.1. protocol, as described previously [14]. The extracted samples were analyzed for the presence of ASFV DNA by qPCR or CSFV RNA by RT-qPCR using the CFX Opus Real-Time PCR System (Bio-Rad, Hercules, CA, USA), essentially as described [27, 28]. For both assays, a positive result was defined as a threshold cycle value (Ct) at which FAM dye emission appeared above background within 42 cycles.

#### Genomic and mitochondrial genome detection

For host DNA detection, the level of the *Sus scrofa* cytoskeletal β-actin gene in the samples was determined using the ACTB-F and ACTB-R primers as described [27], while the level of the *Sus scrofa* mitochondrial cytochrome b gene was determined using an assay developed by Forth [29]. The qPCRs were performed using the CFX Opus Real-Time PCR System (Bio-Rad). A positive result was defined as a FAM (mitochondrial cytochrome b gene) or HEX dye emission signal (β-actin gene) appearing above background within 42 cycles (reported as Ct-values).

#### Genome copy number determination

Absolute quantification was used to determine the number of genome copies per ml by reference to standard curves based on an endpoint dilution series of the ASFV pVP72 plasmid [23] or generated from endpoint dilutions of artificially synthesized double stranded cDNA (dsDNA) (from gBlock, Integrated DNA Technologies, Coralville, IA, USA). The chemically synthesized double stranded cDNA corresponded to the nt 631-763 of the *Sus scrofa* actin mRNA (GenBank: KU672525.1) and the nt 563-856 of the *Sus scrofa* isolate CRB3254 cytochrome b (CYB) gene (GenBank: KY236028.1).

#### Lactate dehydrogenase (LDH) assays

LDH activity was quantified in the serum samples using a lactate dehydrogenase activity assay kit (catalog number MAK066, Sigma-Aldrich, St. Louis, MO, USA). The assays were performed according to the manufacturer’s instructions using serum samples diluted 1:10 in 1x Dulbecco’s phosphate buffered saline (1× DPBS) (Gibco Thermo Fischer Scientific, Waltham, MA, USA) in order to ensure that all measurements were within the linear range of the assay. Measurements were made using a SunriseTM absorbance microplate reader (Tecan, Männedorf, Switzerland). Results were calculated according to the manufacturer’s instructions and presented as milliUnits (mU)/mL.

#### DNA fragment size determination

For DNA size fragment size determination, DNA was isolated from serum samples manually using the TRIzolTM Reagent (Thermo Fischer Scientific) according to the manufacturer’s instructions with minor modifications. Phase separation was achieved using 1- bromo-2-chloropropane (Thermo Fischer Scientific), and following ethanol precipitation, the DNA was resuspended in TE buffer (Thermo Fischer Scientific). Following this DNA purification, AMPure XP beads (Beckman Coulter Life Sciences, Indianapolis, IN, USA) were used to increase DNA concentration and purity. The DNA was then analyzed using the Genomic DNA ScreenTape (Agilent, Santa Clara, CA, USA) on a 4200 TapeStation system (Agilent).

#### Data presentation

Data analysis and presentation were performed using GraphPad prism 9.0 (GraphPad Software, Boston, MA, USA)

## Results

### Characterization of ASFV infections in pigs - Experiment A

#### Assessment of body temperature, clinical signs and virus replication

A group of 4 pigs (numbered 13, 14, 15 and 20) had been inoculated, using the intranasal route, with the highly virulent genotype II ASFV/POL/2015/Podlaskie, as described in Materials and Methods. EDTA-blood samples were collected from each pig prior to inoculation (day 0) and at 3, 5, 6 and 7 days post inoculation (dpi). These whole blood samples, from each sampling day, were assayed for the presence of ASFV DNA by qPCR; note that results are presented in the graphs as 42-Ct values (see Figure 1A). As expected, no ASFV DNA was present at 0 dpi. Low levels of ASFV DNA were detected at 3 dpi in 3 of the 4 pigs (as observed previously using this virus isolate, [23]) and much higher levels were observed at 5, 6 and 7 dpi. Most of the animals were euthanized at 7 dpi but pig 13 was euthanized on day 6 due to severe clinical disease and thus could not be sampled on day 7.

**Figure 1.**
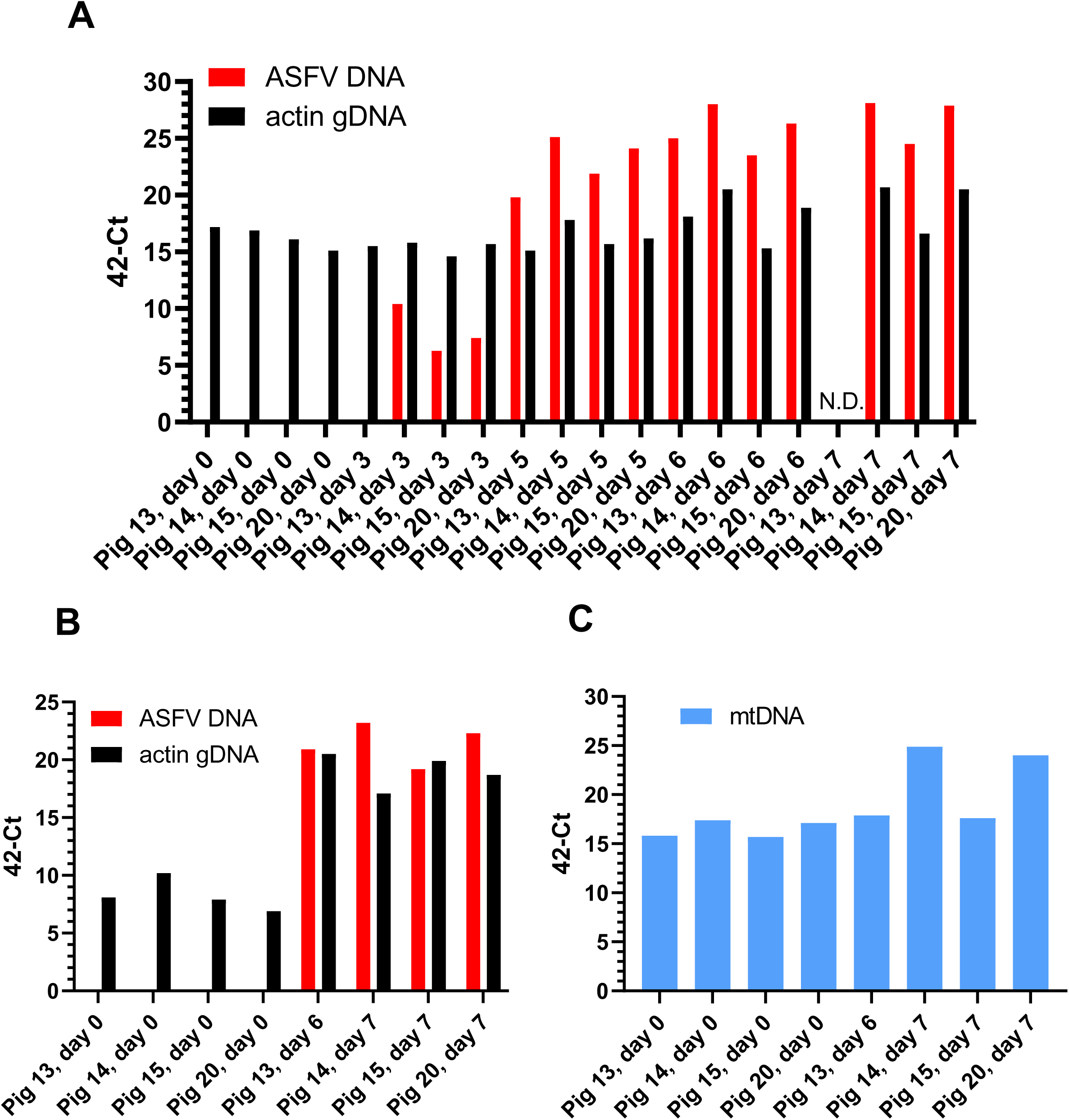
Detection of ASFV genomes and cellular DNAs in whole blood and serum from ASFV-inoculated pigs in Experiment. **A.** Pigs were inoculated by the intranasal route on day 0 and blood samples were collected on the indicated days. Following extraction of nucleic acids the samples were assayed by qPCR. Panel A. Nucleic acids from whole blood samples were assayed in a duplex assay for the presence of the cellular β-actin gene (as a marker for DNA extraction) and for ASFV DNA as indicated. N.D. indicates not determined as pig 13 was euthanized at 6 dpi. Panel B. Nucleic acids extracted from serum samples collected on the indicated days were assayed, as for panel A, for the cellular β-actin gene within genomic DNA (gDNA) and ASFV DNA as indicated. Panel C. Nucleic acids from serum, as used in panel B, were assayed for the presence of mtDNA (targeting the mitochondrial cytochrome b gene). All results are presented as 42-Ct. When no Ct was obtained after 42 cycles, the samples were given a value of 42.

The body temperature and a clinical score (determined as described in Materials and Methods) for each pig were also recorded on each of the sampling days (see Figure 2). A marked increase in body temperature (Figure 2A) was apparent in each pig at 5 dpi and various clinical signs (e.g. lethargy and anorexia) also began to appear at this time (Figure 2B) and developed further to give the highest clinical scores at 6 and 7 dpi. Prior to euthanasia, reddening of the skin and neurological signs (unsteady walk and sometimes convulsions) were observed. As indicated above, pig 13 had to be euthanized at 6 dpi. As may be expected, the increased body temperatures and elevated clinical scores coincided with the large increase in the presence of ASFV DNA in the blood (Figure 1A) essentially from 5 dpi.

**Figure 2.**
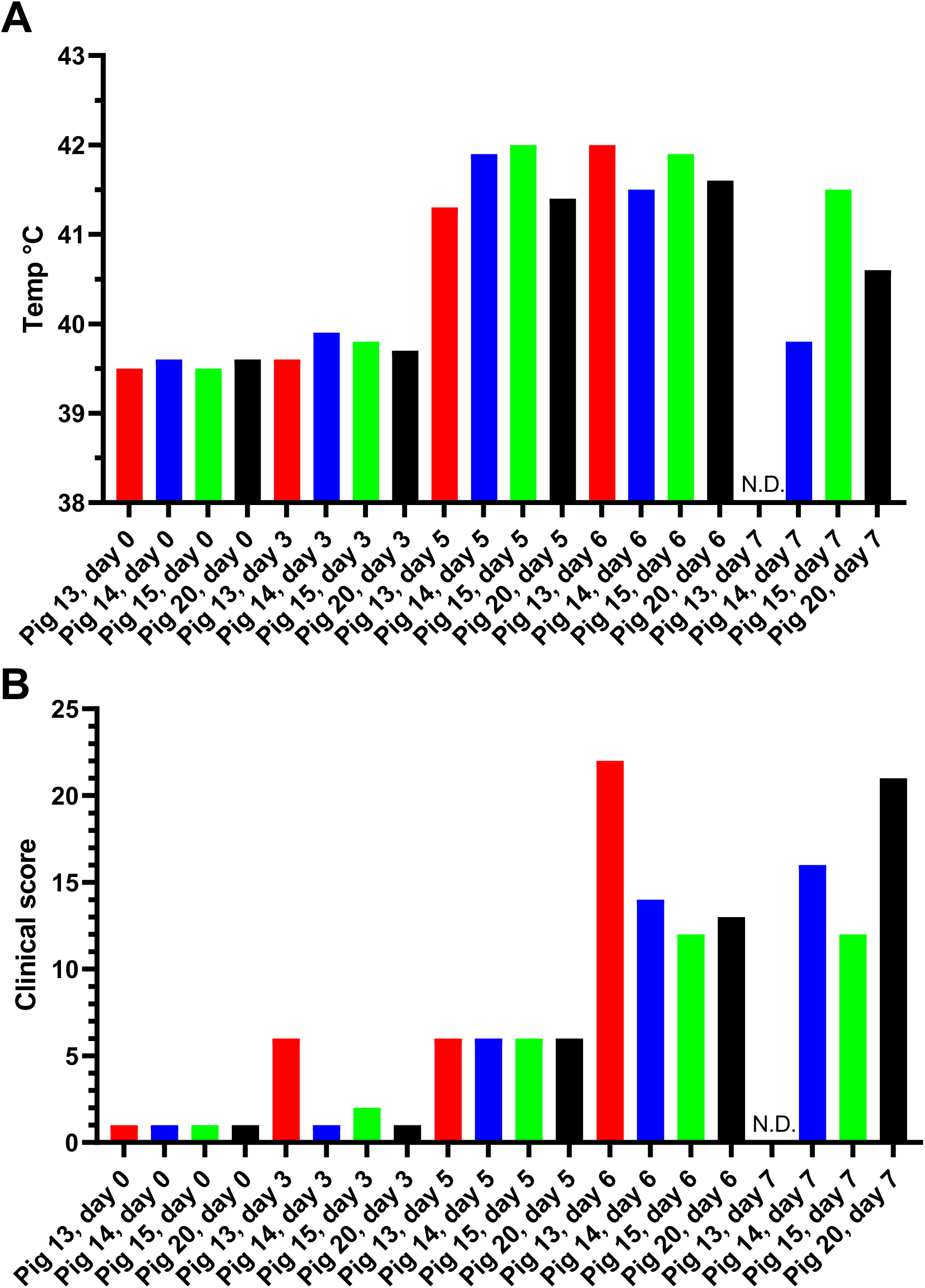
Rectal temperatures and clinical scores in pigs inoculated with ASFV (Podlaskie) in Experiment. **A.** The indicated pigs were inoculated with ASFV Podlaskie, as in Figure 1, the rectal temperatures (panel A) and clinical scores (panel B), were assessed as described in Materials and Methods and the results, from the same days as blood sampling occurred (as shown in Figure 1), are shown. N.D. indicates “not determined” as the pig had been euthanized at 6 dpi.

To ensure that the DNA extractions and qPCR assays were functional, the ASFV DNA was assayed as part of a duplex assay including primers and probes that also detected the gene encoding the cytoskeletal β-actin (part of the genomic DNA (gDNA)) [30]. As expected, high levels of this β-actin gene were detected in the whole blood samples that included both the nucleated white blood cells (e.g. PBMCs) and the enucleated erythrocytes that were collected from each pig on each sampling day (Figure 1A). However, it was noticed that markedly elevated levels of β-actin gDNA were detected in the blood from 2 of the 3 pig samples taken at 7 dpi (i.e. from pigs 14 and 20) when very high levels of ASFV DNA were also detected. The higher levels of β-actin gDNA in the blood were also detected in these two pigs at 6 dpi (Figure 1A).

It seemed possible that the elevated signals for β-actin gDNA at 6 and 7 dpi (Figure 1A) resulted from the destruction of ASFV-infected blood cells that could result in the release of cellular genomic DNA into the blood. To test for the release of cellular DNA into the blood, serum samples (lacking all blood cells) that had been prepared, from the same group of animals, using unstabilized blood samples, collected prior to infection and at euthanasia (at 6 or 7 dpi) were also assayed for the presence of β-actin gDNA. At 0 dpi, the levels of β-actin gDNA in the serum were low (Ct values 32.1-34.8) although readily detectable, see Figure 1B, but at euthanasia, the level of gDNA was much higher (Ct values 21.1-25.2), i.e. a difference of about 10 cycles (ca. 1000-fold increase, as 210=1024). Consistent with the assays using whole blood, no ASFV DNA was detected in the sera at 0 dpi (no Ct value) but very high levels of ASFV DNA were present in the sera at euthanasia (Ct values 19.3 – 23.3) (Figure 1B).

The levels of mitochondrial DNA (mtDNA) were also assessed in the serum samples (Figure 1C) using a qPCR that targeted the mitochondrial cytochrome b gene (see Materials and Methods). Quite high levels of mtDNA were found in the serum of uninfected pigs (Ct values of about 24), however there was a marked increase in the level of mtDNA (about 6 Ct lower at around 18, which represents a change of about 26 = 64-fold) in the serum from pigs 14 and 20 at 7 dpi. These two pigs also had the highest level of ASFV DNA in their serum at that time (Figure 1B).

### ASFV-infection of pigs- Experiment B

To confirm these results, serum samples from a similar but separate experiment, termed here Experiment B, that was performed in 2020 [23, 25] were analyzed. In this experiment, sera had been collected throughout the time course of infection, from 3 separate groups of ASFV-infected pigs, and were tested here for the presence of ASFV DNA, β-actin gDNA and mtDNA. The Ct values from this experiment are presented within the Supplementary Figure S1. To enable easy comparison of the levels of ASFV DNA, mtDNA and gDNA in the serum samples from pigs 1-12, the Ct values obtained (and presented as 42- Ct values in Supplementary Figure S1) were converted, by reference to standard curves, to genome copy numbers/ml and are shown in Figure 3A, B, C.

**Figure 3.**
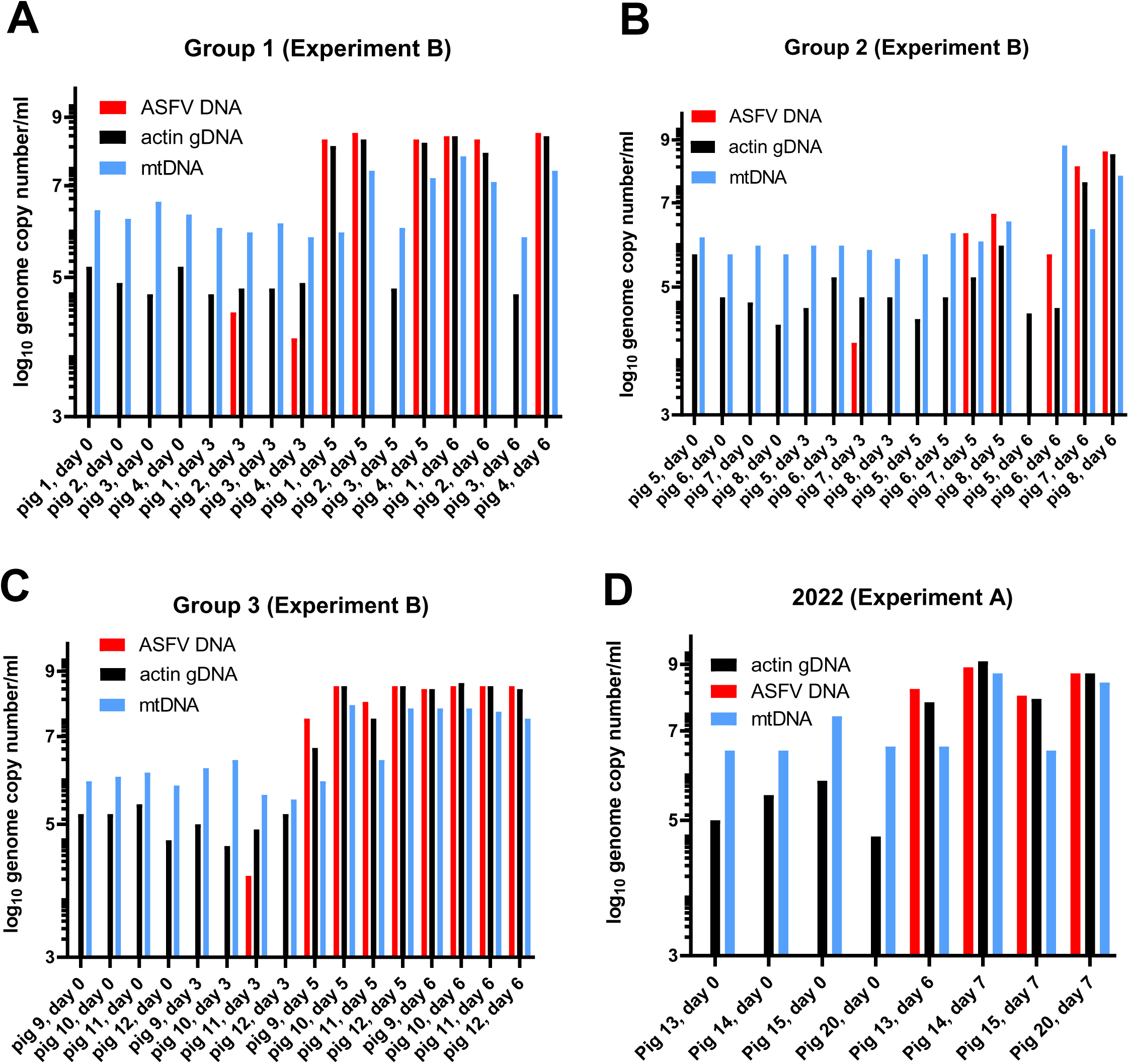
Absolute levels of ASFV DNA, β-actin gDNA and mtDNA in serum from ASFV-inoculated pigs in Experiment B. Nucleic acids isolated from serum samples collected from three separate groups of ASFV-inoculated pigs from Experiment B (as described by Olesen et al.[23]) were assayed for the presence of ASFV DNA, β-actin gDNA and mtDNA and absolute copy numbers/ml were determined from standard curves (panels A, B, C for pigs in Groups 1, 2 and 3 respectively). Note, pigs 3 and 5 did not show any sign of infection by the virus (the temperatures remained normal and there were no clinical signs of disease). The Ct values are shown in Supplementary Figure S1. Data from Experiment A (as shown in Figure 1B, C) were also converted to absolute genome copy numbers and are shown in panel D.

Prior to inoculation at 0 dpi, no ASFV DNA was present in the animals (Figure 3A, B, C, Supplementary Figure S1), however at 3 dpi low levels of the ASFV genome were present in four animals (mean value in these 4 animals (pigs 2, 4, 7 and 11) was ca. 1.5 x 104 ASFV genomes/ml). By 5 dpi, the level of ASFV DNA in serum had increased dramatically to between 106.5 to108.5 genomes/ml (for pigs 1, 2, 4 and 10-12, the mean value was 1.95 x 108 genomes/ml) and, when euthanized at 6 dpi, the level of viral DNA remained very high at up to 108.6 genome copies/ml (the mean value for pigs 1, 2, 4, 7-12 was 2.7 x 108 genome copies/ml serum).

The level of the β-actin gene, as cfDNA, was very consistent, on 0 dpi, at about 105 genome copies/ml (mean = 1.33 x 105 copies/ml) in the serum of the 12 pigs (Figure 3A, B, C). It was little changed at 3 dpi (mean = 1.43 x 105 copies/ml) but had markedly increased at 5 dpi in the ASFV-infected animals at up to about 108 genome copies /ml (mean for pigs 1, 2, 4, 7-12 = 2.4 x 108 copies/ml) and continued at this high level (mean for pigs 1, 2, 4, 7-12 = 4.4 x 108 genomes copies/ml) at 6 dpi when these infected pigs also had very high levels of ASFV DNA in their blood. Thus, during the time course of ASFV infection, the mean level of gDNA (as measured by the level of the β-actin gene) in the sera had increased by over 3000-fold. Furthermore, there is an apparent correspondance between the accumulation of ASFV DNA in serum and the increased presence of cfDNA, containing the β-actin gDNA.

The level of mtDNA in serum (see Figure 3A, B, C) for these 12 pigs at 0 dpi was about 106 genome copies/ml (mean value = 2.5 x 106 copies/ml), and was similar at 3 dpi (mean value = 1.7 x 106 copies/ml). At 5 dpi, some of the pigs had markedly elevated levels of mtDNA (i.e. pigs 2, 4, 10 and 12), with a level of well over 107 genomes /ml (mean value for these 4 pigs = 7.5 x 107 copies/ml). It is noteworthy that these 4 pigs also had very high levels of ASFV DNA and β-actin gDNA in their serum at this time (Figure 3A, B, C).

Finally, at 6 dpi, most of the pigs that were infected with ASFV also had markedly elevated levels of mtDNA in their serum with nearly 108 genomes/ml (mean value for pigs 1, 2, 4, 7- 12 = 7.5 x 107 copies/ml). Thus, between 0 and 6 dpi, the level of mtDNA in the serum of ASFV–infected pigs increased by about 30-fold. It is noteworthy that, in total, 9 of the pigs had markedly increased levels of mtDNA at 6 dpi but, interestingly, only four of these pigs (pigs 2, 4, 10 and 12) had markedly elevated mtDNA at 5 dpi (Figure 3A, B, C) although some other sera, i.e. from pigs 1, 9 and 11, contained high levels of both ASFV DNA and β-actin gDNA on each of these days (Figure 3A, B, C).

These results are similar, both qualitatively and quantitatively, to those observed in the ASFV-infection experiment A (Figure 1 and Figure 3D).

It appears that the level of mtDNA in serum is increased by ASFV infection but the extent of this change is less marked than the change in β-actin gDNA (cfDNA), due, in part, to the higher background level of mtDNA in serum from uninfected animals. Furthermore, the change in the level of mtDNA occurs later than the change in the level of gDNA and ASFV DNA.

### Release of LDH activity

Lactate dehydrogenase (LDH) is a cytoplasmic enzyme that can be released by cells when tissue damage occurs (e.g. following a heart attack). To assess whether the release of genomic DNA into blood was accompanied by release of LDH, the serum samples from the 12 ASFV-infected pigs from Experiment B, as analyzed in Figure 3A, B, C, were assayed for the presence of LDH. It was found (Figure 4) that an increase in the level of LDH activity was apparent at 5 or 6 dpi in many (but not all) of the pigs. Thus LDH release appeared to be a less sensitive marker of cell damage due to ASFV infection than the release of cfDNA or mtDNA.

**Figure 4.**
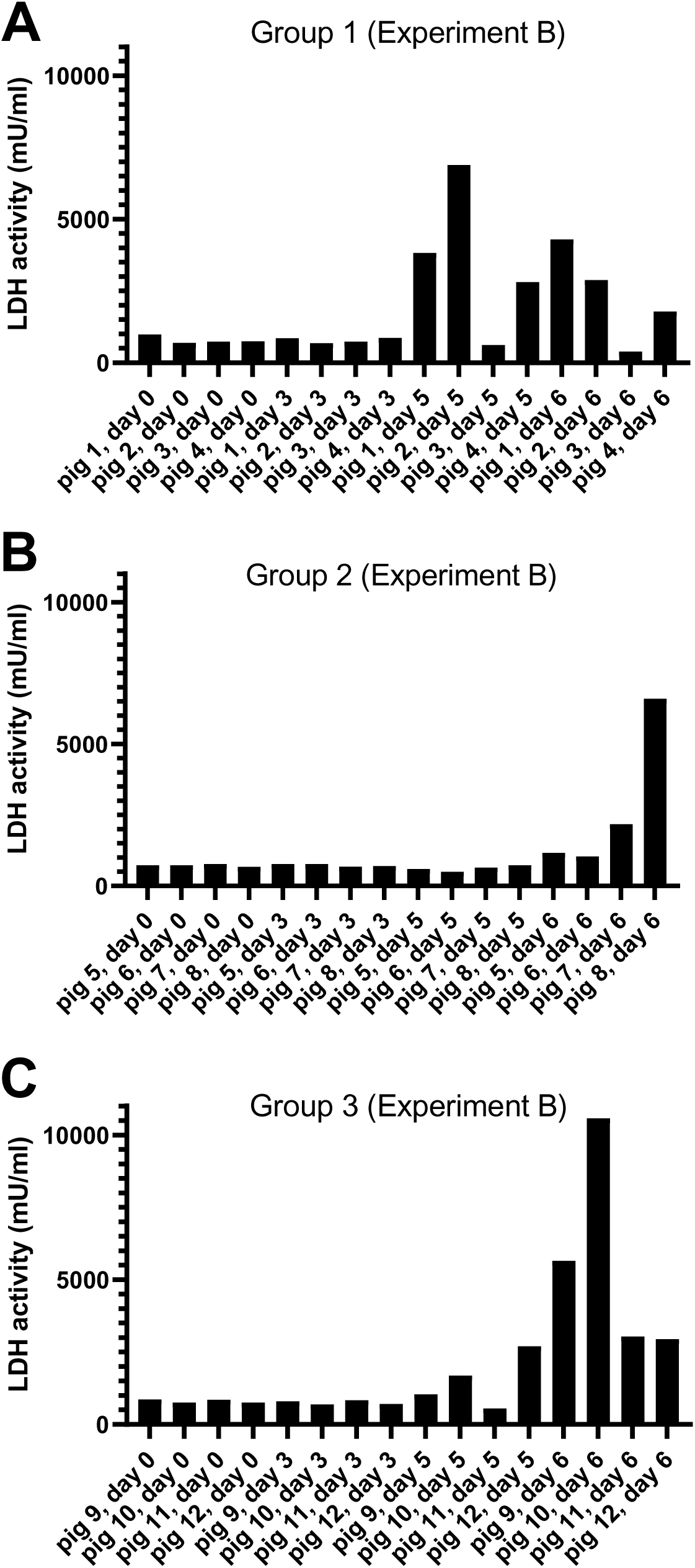
Release of cellular lactate dehydrogenase (LDH) from cells into serum within ASFV-infected pigs (Experiment B). Serum samples, collected on the indicated days, from 12 pigs inoculated with ASFV, in Groups 1-3, as used in Figure 3, were assayed (as described in Materials and Methods) for the presence of the LDH enzyme, a marker for tissue damage.

### Source of cfDNA in ASFV-infected sera

To assess the nature of the cfDNA in the serum of the pigs, selected samples were extracted manually and analyzed to determine the size of the DNA fragments. At 0 and 3 dpi, no DNA fragments were detected in this assay. However, it was found that at 5 or 6 dpi a smear of DNA fragments was present in the extracted samples (Figure 5), these fragments were up to about 1000 bp in length. There was some evidence for specific bands, within the smear, at about 200 bp and 400 bp but there was not a clear ladder of DNA fragments separated by about 180 bp that is indicative of apoptosis.

**Figure 5.**
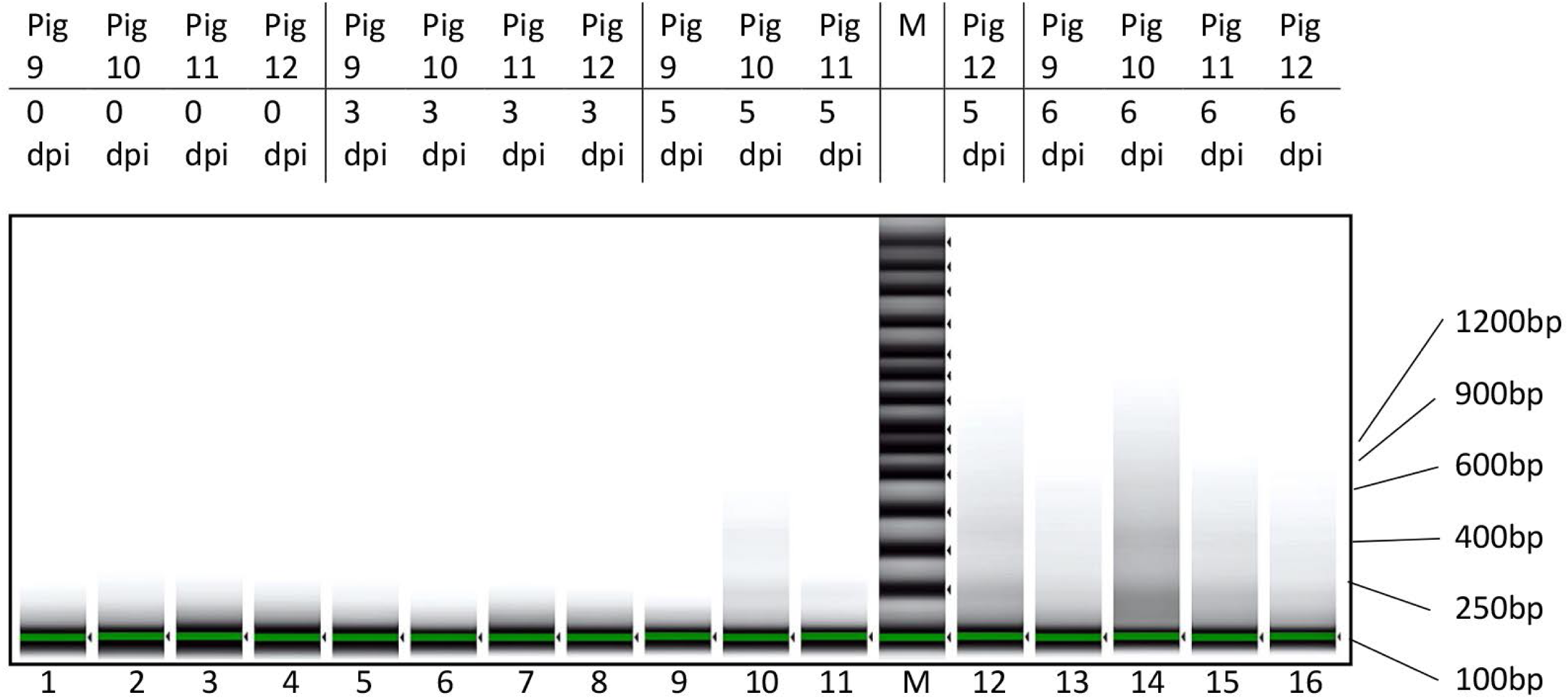
Characterization of DNA fragments in serum of ASFV-infected pigs (Experiment B). Nucleic acids extracted from the serum samples from four different pigs (numbered 9-12) collected on the indicated days post infection (dpi) were purified manually and concentrated (as described in Materials and Methods) prior to analysis on a TapeStation system. The sizes of relevant molecular weight markers are indicated.

### CSFV infection studies

CSFV infection of pigs can cause very similar clinical signs of disease and lymphopenia as observed with ASFV. To determine whether CSFV infection resulted in similar changes in the presence of β-actin gDNA, mtDNA and LDH in the serum of the infected pigs, samples from previously described CSFV-infected pigs [26] were assayed for these markers. Two different strains of CSFV, a low virulence strain, CSFV Bergen, and a highly virulent strain, CSFV Israel, had each been used to infect 3 pigs. Pigs inoculated with CSFV Bergen had viremia at 7 and 10 dpi but 2 of the 3 animals survived and cleared the infection as determined by the loss of CSFV RNA in the serum (Figure 6A), consistent with the earlier results [26]. Pigs inoculated with the CSFV Israel had detectable CSFV RNA in their sera at 4 dpi (Figure 6C) and this increased through to 11 dpi when the animals were euthanized, these results are again consistent with those reported previously [26]. In contrast, the levels of gDNA and mtDNA in the sera remained relatively constant throughout the time course of infection by both strains of CSFV (Figure 6A and 6C). Similarly, there was no apparent change in the level of LDH within the sera of these infected animals throughout the time course of infection (Figure 6B and 6D). Thus, in contrast to the changes in the levels of cfDNA, mtDNA and LDH seen in ASFV-infected pigs (Figures 1-4), there were no marked changes in the levels of these markers within CSFV-infected pigs. It should be noted that the prolonged storage of the samples from the CSFV-infected pigs prior to analysis does not seem to have adversely affected the results. The detection of the viral RNA was consistent with earlier studies on whole blood [26] and the basal levels of LDH were similar to those observed in the recent samples of pig sera obtained prior to inoculation with ASFV (Figure 4).

**Figure 6.**
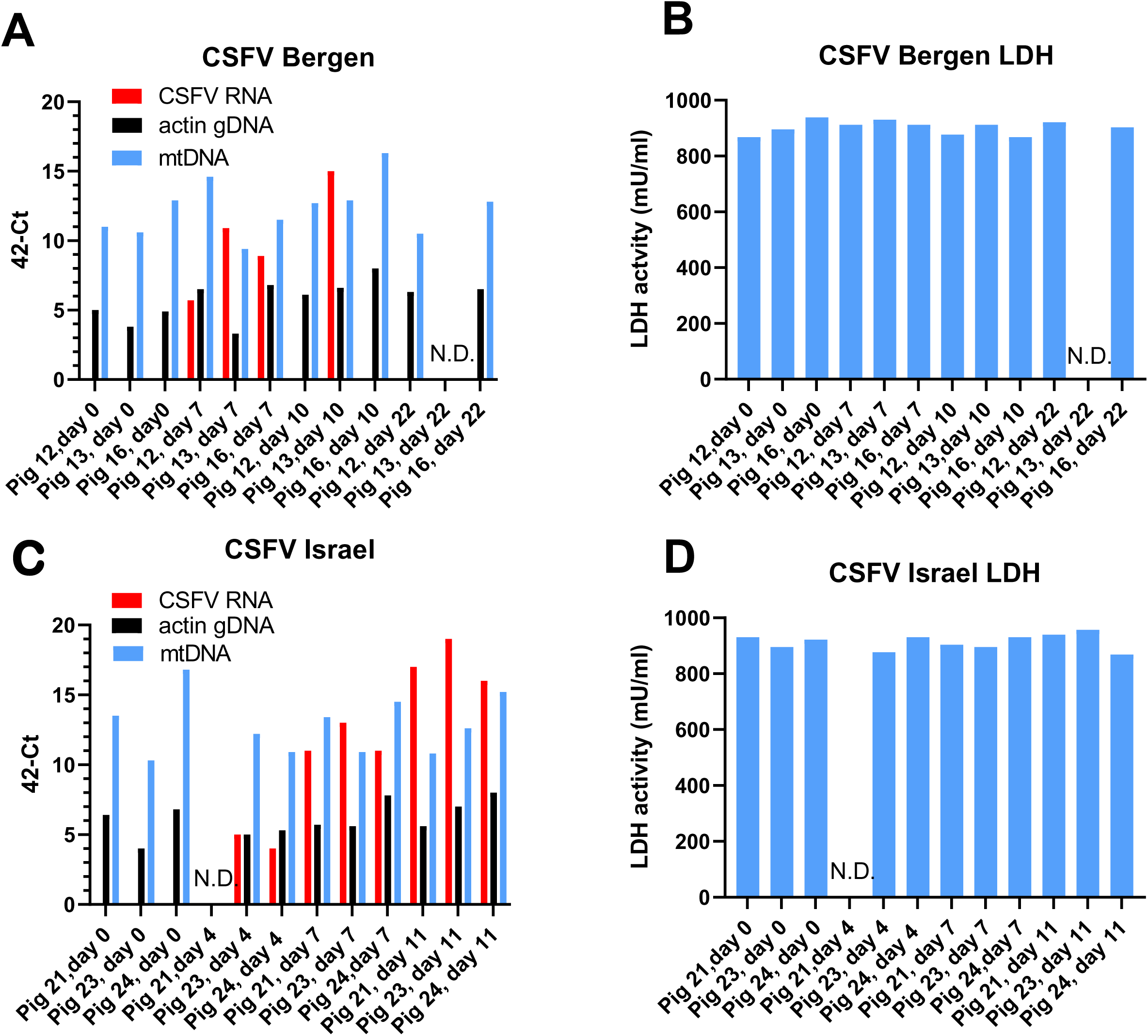
Detection of CSFV genomes, β-actin gDNA and mtDNA in serum from CSFV- inoculated pigs. Pigs were inoculated with the low virulence CSFV Bergen or the highly virulent CSFV Israel (as described by Lohse et al., [26]). Serum samples were collected on the indicated days and stored frozen. Nucleic acids were extracted from these samples and assayed for the presence of CSFV RNA, β-actin gDNA and mtDNA (panels A and C) as described in Materials and Methods. The serum samples were also assayed for the presence of the LDH enzyme (panels B and D). N.D. indicates not determined.

## Discussion

During the course of infection of pigs with a highly virulent ASFV (ASFV/POL/2015/Podlaskie) many changes occur, the animals develop fever and a range of different clinical signs can be apparent. Generally, the animals die within a week of being infected. We have described previously changes in the expression of over 1000 genes in the PBMCs isolated from ASFV-infected pigs [31]. It is well established that major changes occur within the spleen and lymph nodes of ASFV-infected animals [11, 17] and significant loss of B- and T- lymphocytes can occur without these cells being infected themselves, presumably in response to signals received from the infected monocytes [16, 20].

We are unaware of any previous studies that have detected the large (>1000-fold) increase in the level of cfDNA within the sera of ASFV-infected pigs that has been observed here. The parallel detection of ASFV DNA and cfDNA by qPCR may be a convenient way of following the process of infection within the pigs without requiring detection of other virus-specific biomarkers. It seems that the appearance of gDNA in the serum is a much clearer marker of cell death resulting from the ASFV infection than the increased level of mtDNA. The elevated levels of mtDNA in serum were generally detected later in infection, furthermore the mtDNA has a higher baseline signal in the serum of uninfected pigs (see Figures 1 and 3). It had seemed possible that mtDNA would be a better marker for cell death since there are many (hundreds to thousands) mitochondria per cell but there are clearly differences in the way in which the circular mtDNA (ca. 16 kbp) and gDNA (within the nuclei) will be liberated from cells and perhaps the mechanisms for clearance from the circulation will also be distinct. Previous studies [32] have shown that the mean size of mtDNA fragments in plasma is less than 100 bp, this may be because mtDNA, in contrast to gDNA, is not protected by histones and hence does not exist within nucleosomes. The mtDNA normally exists within protein-DNA complexes called nucleoids [33], which contain other DNA binding proteins. The mtDNA is more densely packaged in the nucleoids than the gDNA within nuclei [33]. The reported [32] small size of the fragmented mtDNA may mean that the qPCR assays, in which the targeted mtDNA sequence is 274 bp [29], may be less than optimal. For comparison, the mean size of cfDNA fragments derived from the human genome is ca. 170 bp [32], thus potentially allowing for more efficient detection by qPCR since the targeted sequence is only 114 bp in the β-actin assay [27].

It is likely that nucleated blood cells are among the sources of the fragmented gDNA (cfDNA) [21] but we have not actually demonstrated this due to the expected absence of tissue-specific markers within the short DNA fragments. It is probable that a variety of different cell types (from within the blood itself) and possibly from other tissues (e.g. lymph nodes and spleen) could contribute. From studies in human patients, cfDNA is reported to have only a short half- life in the blood (30 min – 2hrs, [34]). There must be efficient mechanisms to remove the DNA from the blood since apoptosis is a highly regulated mechanism of cell death within an organism with very many cells going through this process on a daily basis [22].

As indicated above, the size of cfDNA is generally small (mean length <200 bp, [32]) and can give an indication of the process of gDNA release. Laddering of the DNA, corresponding to breakage at intervals between nucleosomes (with fragments differing by ca. 200 bp), is consistent with apoptosis whereas necrosis can be expected to yield cfDNA that is more heterogeneous in size [21]. In our studies, we have observed a smear of DNA fragments in samples collected from 5 or 6 dpi (Figure 5) when the level of cfDNA had markedly increased (Figures 1 and 3). There was some evidence for bands at about 200 bp and 400 bp but the pattern did not appear to represent only apoptosis. It may be that the DNA fragments detected are generated by both apoptosis and necrosis. Furthermore, some degradation of the fragments may occur within the serum which could increase the heterogeneity of the fragment sizes.

The release of LDH into the circulation is a convenient marker for cell damage and is widely used for this purpose. The studies presented here do indicate an increase in LDH release into the serum in ASFV-infected animals (Figure 4) and Karalyan et al., [35] have also observed an increase in LDH in sera that resulted from a genotype II ASFV infection in pigs. However, in our studies this change was less consistent than the large increase in the level of cfDNA or the smaller, relative increase in mtDNA identified here. Thus, the production of cfDNA, and to a lesser extent mtDNA, seem to be clear markers for the severe infection within animals produced by the highly virulent genotype II ASFV that is currently circulating in many pig producing countries. The mechanism by which this cfDNA is produced in ASFV-infected pigs is not yet known, however, the absence of such changes in CSFV-infected pigs may suggest a specific role for ASFV-encoded products. Furthermore, the linkage between the virulence of ASFV strains and the release of cfDNA remains to be determined. Potentially, the production of cfDNA may be a useful biomarker for the severity of the infection.

## Acknowledgements

We thank Fie Fisker Brønnum Christiansen, Marie Hornstrup Christensen and Preben Normann for their invaluable technical assistance during this study.

## Author contributions

ASO and CMJ performed the laboratory analyses. GJB drafted the manuscript and prepared the Figures. LL, TBR, AB and GJB supervised the work. All authors reviewed the manuscript.

---

**Supplementary Figure S1.**

**Detection of ASFV DNA, β-actin gDNA and mtDNA in sera from ASFV-inoculated pigs in Experiment B.** Nucleic acids were extracted from serum samples collected on the indicated days from three groups of 4 pigs, (Groups 1 (panel A), Group 2 (panel B) and Group 3 (panel C)) as described in Figure 3. The samples were assayed by qPCR for the presence of ASFV DNA, the cellular β-actin gene (gDNA) and mtDNA as indicated. Results are presented as 42-Ct values. Absolute genome copy numbers derived from these data, using standard curves, are presented in Figure 3A, B and C. As indicated in Figure 3, pigs 3 and 5 showed no indications of becoming infected by ASFV following their inoculation with the virus.

## References

1. Dixon L, Sun H, Roberts H. African swine fever. Antivir Res. 2019; 165: 34–41.

2. Sauter-Louis C, Conraths FJ, Probst C, Blohm U, Schulz K, Sehl J et al. African Swine Fever in Wild Boar in Europe-A Review. Viruses. 2021; 13(9): 1717. doi: 10.3390/v13091717

3. Alonso C, Borca M, Dixon L, Revilla Y, Rodriguez F, Escribano JM, ICTV Report Consortium. ICTV Virus Taxonomy Profile: Asfarviridae. J Gen Virol. 2018; 99(5): 613–614. doi: 10.1099/jgv.0.001049

4. Cackett G, Matelska D, Sýkora M, Portugal R, Malecki M, Bähler J et al. The African Swine Fever Virus Transcriptome. J. Virol. 2020; 94: e00119–20

5. Bastos ADS, Penrith M-L, Crucière C, Edrich JL, Hutchings G, Roger F et al. Genotyping field strains of African swine fever virus by partial p72 gene characterisation. Arch Virol. 2003; 48: 693–706.

6. Dixon LK, Stahl K, Jori F, Vial L, Pfeiffer DU. African swine fever epidemiology and control. Annu Rev Anim Biosci. 2020; 8: 221–246. doi: 10.1146/annurev-animal-021419-08374

7. Ruiz-Saenz J, Diaz A, Bonilla-Aldana DK, Rodríguez-Morales AJ, Martinez-Gutierrez M, Aguilar PV. African swine fever virus: A re-emerging threat to the swine industry and food security in the Americas. Front Microbiol. 2022;13:1011891. doi: 10.3389/fmicb.2022.1011891.

8. EFSA, Ståhl K, Boklund A, Podgórski T, Vergne T, Abrahantes JC et al. Epidemiological analysis of Afrcian swine fever in the European Union during 2022. EFSA J. 2023; 21(5); e08016. 10.2903/j.efsa.2023.8016

9. Mighell E, Ward MP. African Swine Fever spread across Asia, 2018-2019. Transbound Emerg Dis. 2021; 68(5): 2722–2732. doi: 10.1111/tbed.14039.

10. Blome S, Franzke K, Beer M. African swine fever - A review of current knowledge. Virus Res. 2020; 287:198099. doi: 10.1016/j.virusres.2020.198099.

11. Salguero FJ. Comparative Pathology and Pathogenesis of African Swine Fever Infection in Swine. Front Vet Sci. 2020; 7:282. doi: 10.3389/fvets.2020.00282

12. Gallardo C, Soler A, Nieto R, Cano C, Pelayo V, Sánchez MA et al. Experimental Infection of Domestic Pigs with African Swine Fever Virus Lithuania 2014 Genotype II Field Isolate. Transbound Emerg Dis. 2017; 64: 300–304. 10.1111/tbed.12346

13. Nurmoja I, Mõtus K, Kristian M, Niine T, Schulz K, Depner K et al. Epidemiological analysis of the 2015–2017 African swine fever outbreaks in Estonia, Prev Vet Med. 2020: 181: 1044556. 10.1016/j.prevetmed.2018.10.001.

14. Olesen AS, Lohse L, Boklund A, Halasa T, Gallardo C, Pejsak Z et al. Transmission of African swine fever virus from infected pigs by direct contact and aerosol routes. Vet Microbiol. 2017; 211: 92–102. 10.1016/j.vetmic.2017.10.004

15. Guinat C, Reis AL, Netherton CL, Goatley L, Pfeiffer DU, Dixon L. Dynamics of African swine fever virus shedding and excretion in domestic pigs infected by intramuscular inoculation and contact transmission. Vet Res. 2014; 45: 93. 10.1186/s13567-014-0093-8

16. Blome S, Gabriel C, Beer M. Pathogenesis of African swine fever in domestic pigs and European wild boar. Virus Res. 2013; 173(1): 122–130. doi: 10.1016/j.virusres.2012.10.026.

17. Pikalo J, Zani L, Hühr J, Beer M, Blome S. Pathogenesis of African swine fever in domestic pigs and European wild boar – Lessons learned from recent animal trials. Virus Res. 2019; 271: 197614. 10.1016/j.virusres.2019.04.001

18. Oura CA, Powell PP, Parkhouse RM. African swine fever: a disease characterized by apoptosis. J Gen Virol. 1998; 79 (Pt 6):1427–1438. doi: 10.1099/0022-1317-79-6-1427.

19. Ramiro-Ibanez F, Ortega A, Brun A, Escribano JM, Alonso C. Apoptosis: a mechanism of cell killing and lymphoid organ impairment during acute African swine fever virus infection. J Gen Virol. 1996; 77:2209–2219

20. Ramiro-Ibanez F, Ortega A, Ruiz-Gonzalvo F, Escribano JM, Alonso C. Modulation of immune cell populations and activation markers in the pathogenesis of African swine fever virus infection. Virus Res. 1997; 47 (1): 31–40.

21. Celec P, Vlková B, Lauková L, Bábíčková J, Boor P. Cell-free DNA: the role in pathophysiology and as a biomarker in kidney diseases. Expert Rev Mol Med. 2018; 20:e1. doi: 10.1017/erm.2017.12.

22. Kakarla R, Hur J, Kim YJ, Kim J, Chwae YJ. Apoptotic cell-derived exosomes: messages from dying cells. Exp Mol Med. 2020; 52(1):1–6. doi: 10.1038/s12276-019-0362-8.

23. Olesen AS, Kodama M, Lohse L, Accensi F, Rasmussen TB, Lazov CM et al. Identification of African Swine Fever Virus Transcription within Peripheral Blood Mononuclear Cells of Acutely Infected Pigs. Viruses. 2021 Nov 22;13(11):2333. doi: 10.3390/v13112333.

24. Olesen AS, Lazov CM, Lecocq A, Accensi F, Jensen AB, Lohse L et al. Uptake and Survival of African Swine Fever Virus in Mealworm (*Tenebrio molitor*) and Black Soldier Fly (*Hermetia illucens*) Larvae. Pathogens 2023; 12: 47. Doi: 10.3390/pathogens12010047

25. Olesen AS, Lohse L, Accensi F, Goldswain H, Belsham GJ, Bøtner A et al. Inefficient transmission of African swine fever virus to sentinel pigs from environmental contamination under experimental conditions. Transbound Emerg Dis. 2023; (in revision).

26. Lohse L, Nielsen J, Uttenthal Å. Early pathogenesis of classical swine fever virus (CSFV) strains in Danish pigs. Vet. Microbiol. 2012; 159:327–336.

27. Tignon M, Gallardo C, Iscaro C, Hutet E, Van der Stede Y, Kolbasov D et al. Development and inter-laboratory validation study of an improved new real-time PCR assay with internal control for detection and laboratory diagnosis of African swine fever virus. J Virol Methods. 2011; 178: 161–170. 10.1016/j.jviromet.2011.09.007

28. Hoffmann B, Beer M, Schelp C, Schirrmeier H, Depner K. Validation of real-time RT-PCR assay for sensitive and specific detection of classical swine fever. J Virol Methods. 2005; 130: 36–44.

29. Forth JH. Standardisierung Eines Nicht-Invasiven Beprobungssystems zur Infektionsüberwachung bei Wildschweinen (Sus scrofa); Ernst-Moritz-Arndt- Universität Greifswald, Mathematisch-Naturwissenschaftliche Fakultät: Greifswald, Germany. 2015. Available online: https://www.openagrar.de/receive/openagrar_mods_00012495 (accessed on 13 July 2023).

30. Dahlén A, Mertens F, Mandahl N, Panagopoulos I. ACTB (Actin, beta). Atlas Cytogenet Oncol Haematol. 2005; 9(2):129–130.

31. Olesen AS, Kodama M, Skovgaard K, Møbjerg A, Lohse L, Limborg MT et al. Influence of African Swine Fever Virus on Host Gene Transcription within Peripheral Blood Mononuclear Cells from Infected Pigs. Viruses. 2022;14(10):2147. doi: 10.3390/v14102147.

32. Phung Q, Lin MJ, Xie H, Greninger AL. Fragment Size-Based Enrichment of Viral Sequences in Plasma Cell-Free DNA. J Mol Diagn. 2022;24(5):476–484. doi: 10.1016/j.jmoldx.2022.01.007.

33. Bogenhagen DF. Mitochondrial DNA nucleoid structure. BBA- Gene Regulatory Mechanisms. 2012; 1819: 914–920. 10.1016/j.bbagrm.2011.11.005

34. Sherwood K, Weimer ET. Characteristics, properties, and potential applications of circulating cell-free DNA in clinical diagnostics: a focus on transplantation. J Immunol Meth. 2018; 463: 27–38. 10.1016/j.jim.2018.09.011.

35. Karalyan Z, Zakaryan H, Arakelova E, Aivazyan V, Tatoyan M, Kotsinyan A et al. Evidence of hemolysis in pigs infected with highly virulent African swine fever virus. Veterinary World. 2016; 9(12): 1413–1419

