## Supplementary figures and images for "Release of fragmented host, cell-free, genomic DNA into the circulation of pigs during infection by virulent African swine fever virus"

### Supplementary Figure

#### Supplementary Figure S1

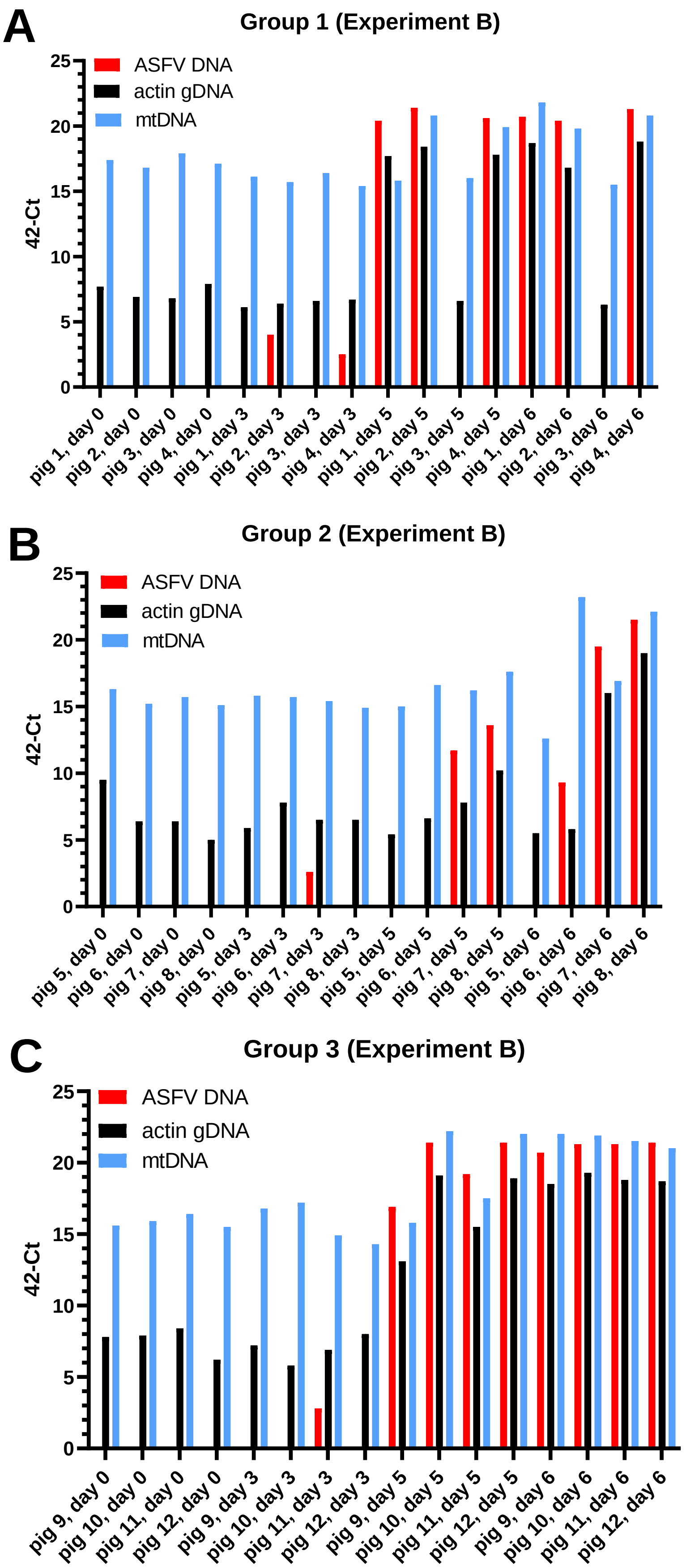
